# Can higher-order interactions resolve the species coexistence paradox?

**DOI:** 10.1101/2023.06.19.545649

**Authors:** Theo Gibbs, Gabriel Gellner, Simon A. Levin, Kevin S. McCann, Alan Hastings, Jonathan M. Levine

## Abstract

Most ecological models are based on the assumption that species interact in pairs. Emerging in diverse communities, however, are higher-order interactions, in which two or more species jointly impact the growth of a third species. A pitfall of the pairwise approach is that it misses the higher-order interactions potentially responsible for maintaining diversity in nature. Here, we explore how well higher-order interactions enable coexistence when pairwise interactions are insufficient to do so. Specifically, we explore the stability properties of systems where higher-order interactions guarantee that a specified set of abundances are a feasible equilibrium of the dynamics. Even these higher-order interactions do not necessarily produce robust coexistence. We find that facilitative higher-order interactions that counter pairwise competitive interactions are less likely to generate stable equilibria than competitive interactions countering pairwise facilitation. In both cases the system is more likely to be stable when the pairwise interactions are weak, and when the equilibrium abundances are near those produced by the pairwise interactions. Last, we show that correlations between the pairwise and higher-order interactions permit robust coexistence in diverse systems. Our work reveals the challenges in generating stable coexistence through higher-order interactions, but also uncovers higher-order interaction patterns that can enable diversity.

## Introduction

Understanding how species stably coexist in diverse communities is one of the central challenges in ecology [1, 2, 3]. Most attempts to solve this problem have assumed that species interact in a pairwise fashion [4], and thus experiments on every pair of species in a community should reveal the complete interaction network [5]. Unfortunately, patterns in this network are rarely sufficient to explain the coexistence observed in nature [6, 7]. One fascinating explanation for this mismatch is that interactions only possible with more than two species in a community are precisely those that sustain natural diversity. Interactions between three or more species, called higher-order interactions, have been studied for decades in ecology [8, 9, 10, 11, 12, 13], partly because they may contribute to coexistence [5, 14, 15, 16, 17, 18], yet empirical and theoretical tests of this expectation are rare [5].

Of the few theoretical studies, those modeling specific biological scenarios have sometimes shown that higher-order interactions can indeed permit widespread coexistence [15, 16]. For example, antibiotic-degrading bacteria can protect antibiotic-sensitive bacteria from the toxins secreted by other bacteria, thus enabling coexistence [15, 19]. More recent theory, however, incorporating higher-order interactions into a general model of interacting species, has shown that randomly assigned higher-order interactions do not increase the likelihood of coexistence any more than randomly sampled pairwise interactions [20]. In short, although higher-order interactions can stabilize coexistence in some specific biological situations, they do not do so generically. As a result, understanding which types of higher-order interaction networks can stabilize coexistence is an important open question. Here, we bridge the apparent gap between results from specific and generic models by determining when higher-order interactions can overcome exclusion caused by pairwise interactions.

We do so by considering the stability properties of systems in which higher-order interactions have been specifically chosen to generate a feasible coexistence equilibrium with given species abundances, even when the pairwise interactions alone would generate competitive exclusion. We then ask if higher-order interactions were to explain the coexistence of the entire community at given abundances, what are the necessary patterns among these interactions? We answer this question with a theoretical model using a method originally developed to study metabolic networks [21], but recently applied to ecology [22]. Specifically, given pairwise interactions and species abundances, we first determine the higher-order interactions necessary to ensure an equilibrium of the dynamics at those abundances. Interestingly, these equilibria are not always stable to small perturbations, meaning that even seemingly optimal higher-order interactions do not necessarily produce robust coexistence. In this context, we address three questions about when and how higher-order interactions can produce stable coexistence: 1) How does the sign and strength of the pairwise interactions influence the stability of coexistence enabled by higher-order interactions? 2) How sensitive is the stability of this coexistence to species’ equilibrium abundances? 3) Can correlations between the pairwise and higher-order interactions ensure stable coexistence?

## Results

### Model form, parameterization and overall approach

We study our research questions using an extension of the classic Lotka-Volterra model that includes higher-order interactions among three species. In a community of *S* species, the dynamics of species *i* are given by

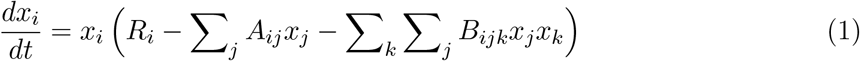

where *R*_*i*_ is the intrinsic growth rate of species *i, A*_*ii*_ measures the strength of self-regulation of species *i*, and *A*_*ij*_ describes the pairwise impact that species *j* has on species *i*’s growth (Fig. 1A). We model the higher-order effect of species *j* and species *k* on species *i* as a scalar on the product of the abundances *x*_*j*_ and *x*_*k*_. The strength of this interaction is given by coefficient *B*_*ijk*_. We only allow higher-order interactions to involve three distinct species, so parameters *B*_*ijk*_ with repeated indices are set to zero. We sampled the inter-specific pairwise interactions from a normal distribution with specified statistics (mean *μ*_*A*_ and variance 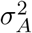), as in many previous approaches [6, 23, 24, 25, 26, 27, 28, 29]. The parameters *μ*_*A*_ and 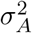 control the strength and variation in the underlying pairwise interactions respectively. Note that the pairwise and higher-order interaction coefficients can be positive (denoting competition) or negative (denoting facilitation).

**Figure 1:**
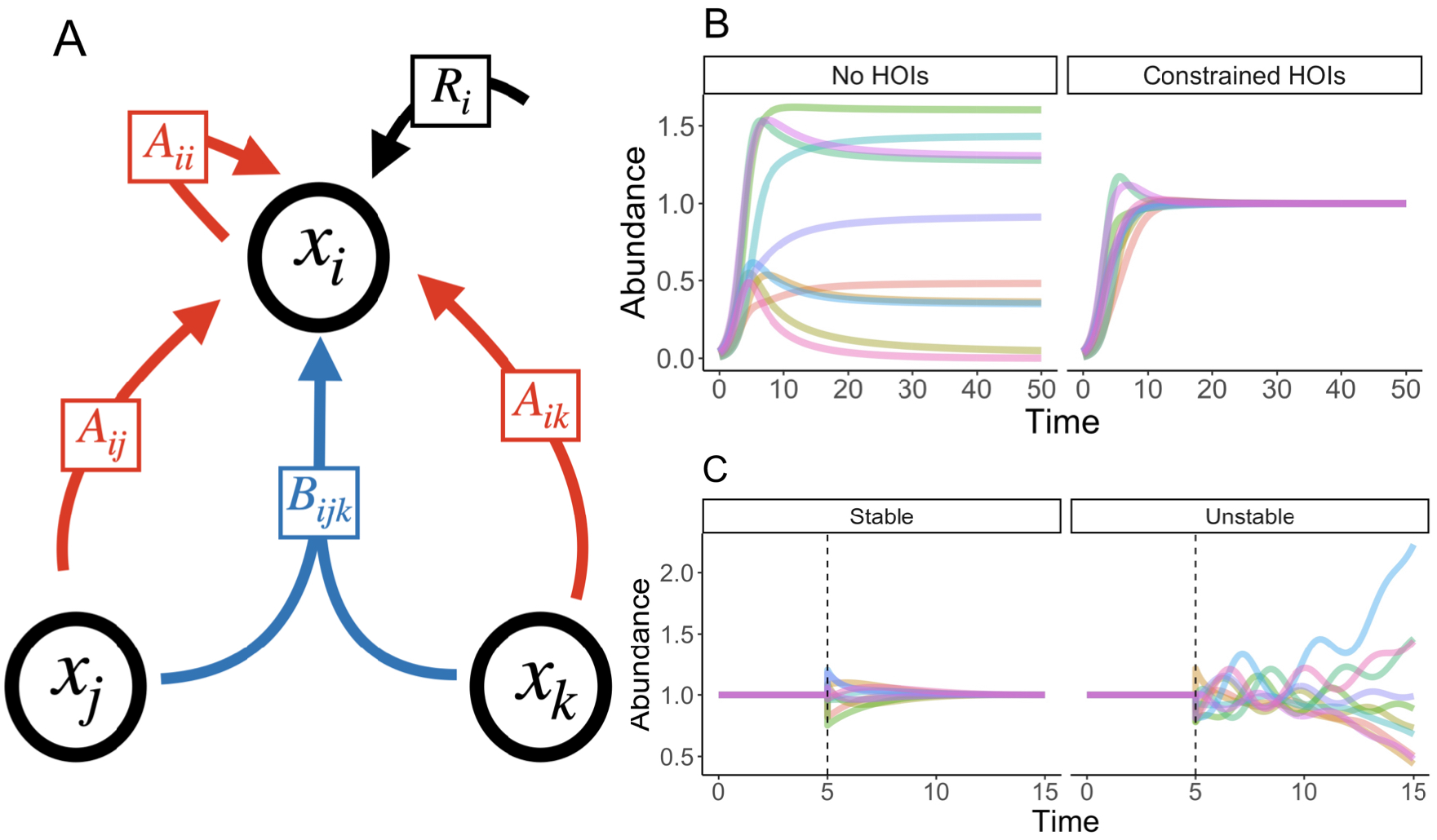
(A) As in Equation 1, species *i* has an intrinsic growth rate *R*_*i*_, experiences intra-specific competition through the parameter *A*_*ii*_ and the pairwise effect of species *j* through the coefficient *A*_*ij*_. The higher-order effect of species *j* and species *k* on species *i* is encoded by the parameter *B*_*ijk*_. (B) Two examples of the dynamics of Equation 1 – one without higher-order interactions (labeled no HOIs) where a single species is excluded and one with the constrained higher-order interactions (labeled constrained HOIs) where all species coexist. (C) We plot simulations of the dynamics of Equation 1 with constrained higher-order interactions when the target abundance equilibrium is stable and unstable. The simulations begin at the target abundance equilibrium until a perturbation (the dashed vertical line) randomly changes the abundances. The resulting dynamics reveal the stability of the equilibrium.

We first asked what type of higher-order interactions would produce a feasible equilibrium – one where all species are at positive abundance [21, 22, 26, 30, 31, 32]. When pairwise interactions would exclude one or more species, we denote the higher-order interactions that reverse these exclusions “constrained higher-order interactions” (see Fig. 1B). Our approach also requires specifying 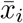, the target abundances of each of the coexisting species. The robustness of our results to the choice of 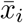 is examined in a later section. Guaranteeing that the target abundances are feasible introduces the following constraints on the higher-order interactions:

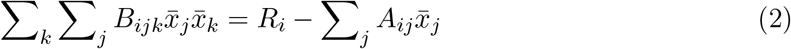

for each *i*. In words, the sum of all higher-order effects on a given species must equal the difference in its intrinsic growth rate and the sum of the pairwise effects it experiences. Note that the constraint is on the sum of the higher-order interactions, not the strength and sign of each interaction itself. Given this, in a diverse community, there are many more higher-order interactions than species, meaning that this constraint problem is severely under-determined as many different choices of the higher-order interactions satisfy Equation 2. Therefore, to generate higher-order interaction parameters for our simulations, we sample uniformly from the space of all possible solutions to Equation 2 where the higher-order interactions have the required sign. For example, when the growth rate and pairwise interactions affecting a given species would would cause it to grow above its target abundance equilibrium (ie. the right hand side of Equation 2 is positive), then all the higher-order interactions affecting that species must be competitive. In the Methods, we lay out the sampling procedure in more detail.

Some properties of the higher-order interactions required for coexistence are immediately apparent from the constraint in Equation 2. As the pairwise interactions become more competitive, for example, the higher-order interactions must be more facilitative to maintain the equality. Similarly, increasing the variation in the pairwise interactions makes the sums of the pairwise interaction strengths (ie. 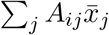) increasingly different across focal species, which, in turn, makes the higher-order interactions more variable across species. Throughout this paper, we characterize coexistence at the equilibrium produced via the constraints in Equation 2, but coexistence at other equilibria or through cyclic or chaotic dynamics is also possible, as examined in the Supplementary Information.

### Question 1: How do the pairwise interaction statistics affect the probability of stability?

**Our first main result is that higher-order interactions that are constrained to produce a feasible equilibrium do not necessarily guarantee coexistence, because these equilibria are not always stable to small perturbations (Fig. 1C)**. Stability instead depends on the statistics of the underlying pairwise interactions. We first delineate these stability rules in the simpler case where the target abundances are equal to the single species carrying capacities of all the species 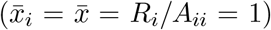. To determine if the equilibrium generated by a given set of interactions is stable, we construct the Jacobian of the model. The Jacobian is a matrix of partial derivatives describing how the abundance of each species changes with a small change in each of the other species and its own species. We evaluate the Jacobian at the target abundance equilibrium and then ask whether or not all of its eigenvalues have a negative real part – the criterion for local stability (see Methods for more details).

**Our second main result is that increasing the overall strength of the pairwise interactions, and concomitantly, the strength of the higher-order interactions needed to counter their effects, decreases the probability of stability (Fig. 2A)**. Regardless of whether the pairwise interactions are competitive or facilitative, as the strength of the pairwise interactions increases, we reach a point at which the probability of stability goes from very high to very low (Fig. 2A). **Moreover, our third main result is that when the system is made feasible by constrained higher-order interactions, facilitative pairwise interactions offset by competitive higher order interactions produce equilibria more likely to be stable than those produced by competitive pairwise interactions offset by facilitative higher order interactions at the same overall magnitude of interaction strength (Fig. 2A)**.

**Figure 2:**
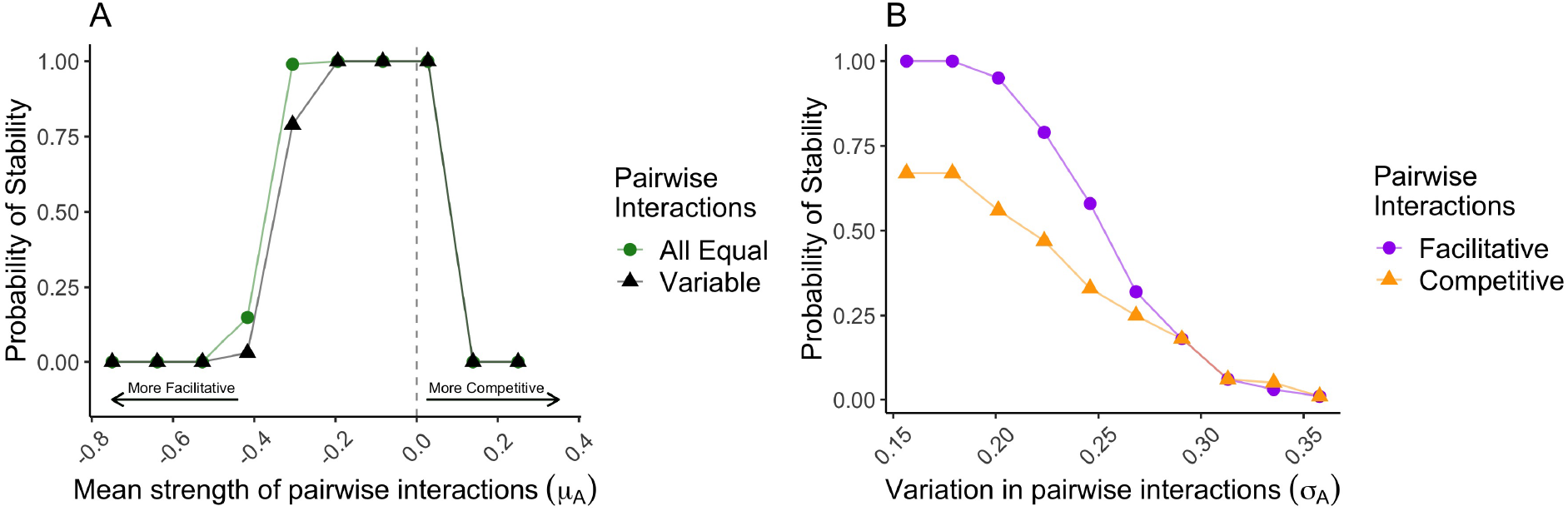
The probability that the target abundance equilibrium is stable as a function of (A) the mean strength of the pairwise interactions or (B) the variation in pairwise interactions computed over 100 replicates. In both panels, the constrained higher-order interactions produce the target abundances as an equilibrium, and we compute whether or not the full system (with both pairwise and higher-order interactions) is stable. In panel (A), the colors and shapes designate different values for the variations in the pairwise interactions – all equal denotes *σ*_*A*_ = 0 and variable denotes *σ*_*A*_ = 0.1. The dashed vertical line displays where the mean interaction strength is zero, and therefore not competitive or facilitative. In panel (B), the colors and shapes designate whether or not the underlying pairwise interactions are competitive (*μ*_*A*_ = 0.05) or facilitative (*μ*_*A*_ = −0.05). In both panels, there are 20 species and the target abundances are all 1.

These two results are a natural extension of classic theory on pairwise interactions. When pairwise interactions are competitive, the higher-order interactions must be facilitative to counter them. Moreover, because each higher-order interaction enters into the Jacobian twice (as explained in Methods), the total effect of the facilitative higher-order interactions is twice the strength of the competitive pairwise interactions. This ultimately gives rise to a Jacobian with stability properties identical to a system with only facilitative pairwise interactions (see Methods for the calculation). Following the same logic, facilitative pairwise systems stabilized by competitive higher-order interactions generate a Jacobian with stability properties identical to a system with competitive pairwise effects. This is important because one lesson from previous theory [23] is that facilitative pairwise interactions produce equilibria that are less likely to be stable than competitive pairwise interactions – the very same behavior seen here with facilitative versus competitive higher-order interactions. Moreover, the stronger the pairwise interactions, the stronger the higher-order interactions needed to counterbalance them, which generates stronger off-diagonal elements of the Jacobian – a feature also known to be destabilizing. We can further show that aside from changing the sign of the elements of the Jacobian (by canceling the pairwise effect), there are no additional stability consequences of the higher-order interactions. We know this because whether we satisfy the condition for feasibility in Equation 2 by including higher-order interactions or by changing the intrinsic growth rates generates an equilibrium with the same stability properties (see Fig. D of the Supplementary Information).

Increasing variation in the pairwise interactions also decreases the probability of stability (Fig. 2B) resonating with both classic pairwise theory [6, 23] and more recent higher-order interaction theory [14, 20]. The variability in the pairwise, rather than higher-order, interactions destabilizes the target abundance equilibrium (see the Methods for an analytical calculation, Fig. A of the Supplementary Information for additional numerical evidence and Fig. C of the Supplementary Information for how different sampling procedures modify this result).

### Question 2 : How do the target abundances affect the probability of stability?

One of the main takeaways of our work thus far has been that stronger higher-order interactions are less likely to create stable equilibria, and this is especially true for facilitative higher-order interactions. We next explored the robustness of this result to the choice of abundances. To do so, we specify the pairwise interactions, set all of the target abundances to a given value 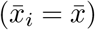 and then compute the probability of stability as a function of this common target abundance (Fig. 3A). Importantly, varying the target abundances changes the required strength of the constrained higher-order interactions, just as varying the pairwise interactions did in our first set of results.

**Figure 3:**
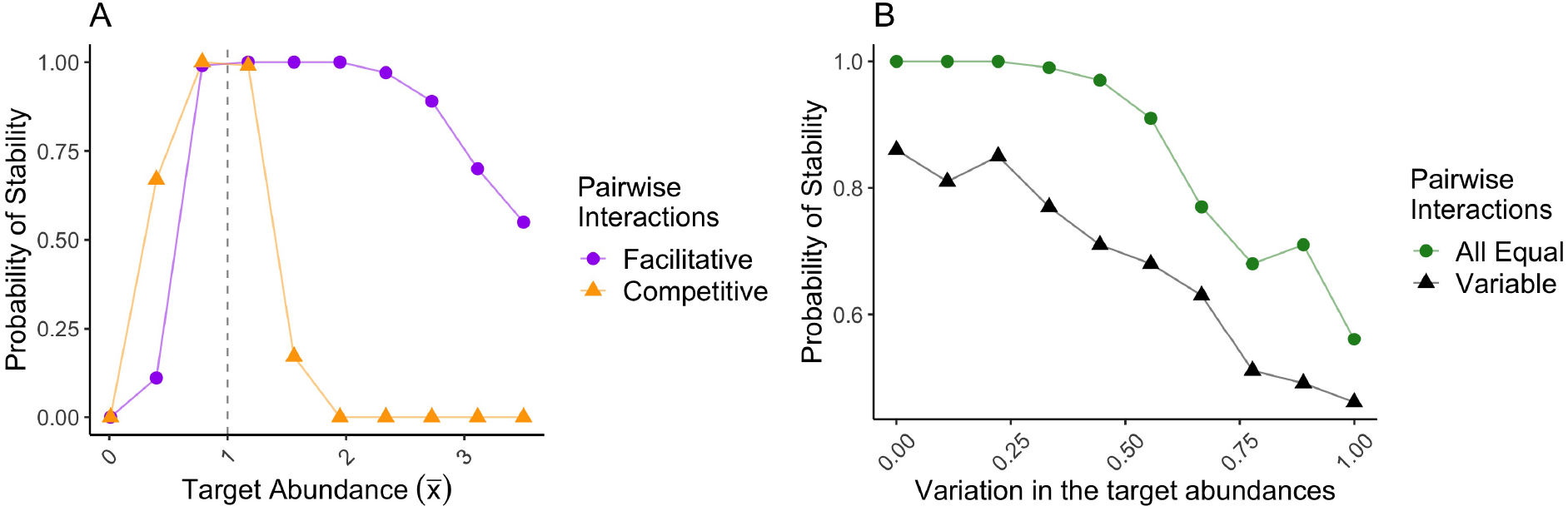
(A) The probability that the target abundance equilibrium is stable as a function of the target abundance equilibrium value 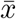. The colors and shapes designate whether or not the mean pairwise interaction strength is facilitative or competitive – specifically, facilitative denotes *μ*_*A*_ = −0.1 and competitive denotes *μ*_*A*_ = 0.1. Constrained higher-order interactions produce the value on the x-axis 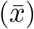 as an equilibrium of the model. The dashed vertical line shows the shared carrying capacity of all the species, which we used as the target abundances in Fig. 2. (B) The probability that the target abundance equilibrium is stable as a function of the variation in the target abundances. The different colors and shapes denote different values for the variation in the underlying pairwise interactions – all equal denotes *σ*_*A*_ = 0 and variable denotes 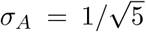. The mean pairwise interaction strength is competitive, specifically *μ*_*A*_ = 0.1. Constrained higher-order interactions produce the target abundances as an equilibrium, and these target abundances become more variable moving right on the x-axis. There are *S* = 5 species and the probabilities are computed over 100 replicates in both panels.

In general, we find that stability increases when the pairwise interactions are of the sign and magnitude required to get the system close to the specified target abundances, and therefore requiring the smallest higher-order effects (regardless of sign). For small target abundances (half the carrying capacity, say), competitive pairwise interactions produce equilibria more likely to be stable than facilitative pairwise interactions (Fig. 3A). This is because competitive pairwise interactions, which get the system close to the low target abundances, require weaker competitive higher-order interactions to get the system all the way there than do facilitative pairwise interactions, which on their own send abundances too high. This finding follows from our second main result that weaker higher-order interactions are more likely to generate a stable equilibrium (see the Supplementary Information for further exploration of this stability landscape). The situation is analogous for target abundances above the carrying capacities, where communities structured by facilitative pairwise interactions are more likely to be stable than those with competitive interactions (Fig. 3A).

We also found that increasing the variance of the probability distribution from which the target abundances are sampled makes these equilibria progressively less likely to be stable (Fig. 3B). As the target abundances become more variable (and more different than would result from the randomly drawn pairwise interactions), the constrained higher-order interactions must also become larger and more variable to produce these abundances as equilibria, eventually leading to instability.

### Question 3: How do correlations between the pairwise and higher-order interactions affect stability?

Thus far, we have explored the stability of equilibria generated by higher-order interactions whose only constraint was their sign and their summed effect on a focal species. Within those constraints, the values of the individual higher-order terms were randomly assigned, meaning that competitors that exerted stronger pairwise effects on a focal species were no more or less likely to be engaged in a stronger higher-order effect on that focal species, or experience such an effect in return. Such relationships between the pairwise and higher-order interactions might not only be biologically justified but may also predictably affect species coexistence. Fortunately, by evaluating the Jacobian, we can identify the specific patterns both in the higher-order interaction network itself and in the correlations between the higher-order and pairwise networks that generate stable equilibria.

To explore how relationships within and between the pairwise and higher-order interaction network influence local stability, we once again focus on the simplified case where the target abundances are the carrying capacities (ie. 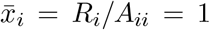. Because the elements of the Jacobian (and local stability) only reflect how the abundances of all species *j* depend on all species *i*, we can sum all higher-order effects of species *j* on species *i* (involving all species *k*) and encode that sum Σ_*k*_(*B*_*ijk*_ + *B*_*ikj*_) into a matrix 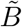 analogous to the pairwise interaction matrix *A*, without any loss of information relevant to local stability. After doing so, the Jacobian of the community evaluated at the target abundance equilibrium is simply the sum of the pairwise and higher-order effects: 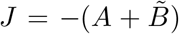 With this framing of the problem, we show that higher-order interactions that give rise to Jacobian entries that are anti-correlated across the diagonal (ie. *J*_*ij*_ ≈ −*J*_*ji*_) promote stability (see Fig. 4 and previous theory for pairwise interactions [23]). Jacobia where this anti-correlation is perfect are called skew-symmetric matrices. Skew-symmetric interaction structures could arise if higher-order interactions that involve species *j* affecting species *i* (in concert with a third species) are of opposite sign to the pairwise effect of species *i* on *j*.

**Figure 4:**
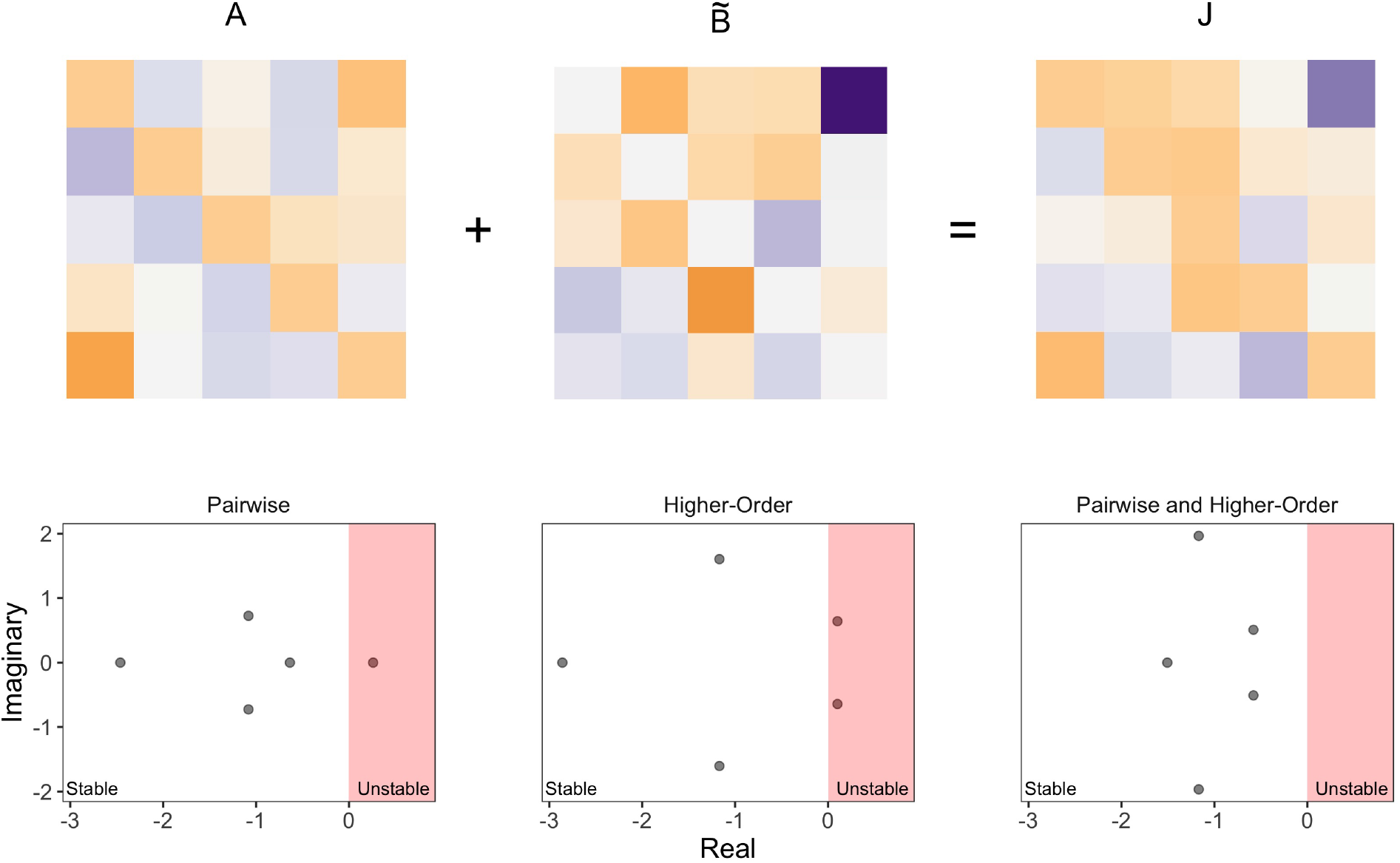
The top row shows three matrices – *A* which is the pairwise interaction matrix, 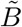 the matrix encoding the higher-order effect of species *j* on species *i* (ie. 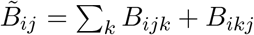) and the Jacobian *J* evaluated at the target abundance equilibrium. More orange (respectively purple) color indicates more competitive (respectively more facilitative) values. The sum of *A* and 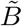 is the Jacobian *J*. Notice that the Jacobian matrix is skew-symmetric – if an entry below the diagonal (the bottom left corner *J*_51_ for example) is strongly competitive, the corresponding entry above the diagonal (the top right corner *J*_15_) is strongly facilitative. Intuitively, the matrix *A* measures all the pairwise interactions in the community, the matrix 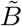 measures all higher-order interactions in the community and the Jacobian *J* combines these two interactions to measure the net effect of each species on every other. The bottom row shows the eigenvalues of the Jacobia of this community when the inter-specific interactions are purely pairwise, purely higher-order or both pairwise and higher-order together. The red shaded region displays all eigenvalues with positive real part. The equilibria are stable when all the eigenvalues in this bottom row have negative real part (in other words, none of the eigenvalues lie in the red shaded region). The communities with only pairwise or only higher-order interactions are unstable, but together they stabilize the community because the Jacobian is approximately skew-symmetric. There are *S* = 5 species, 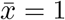, *μ*_*A*_ = 0 and *σ*_*A*_ = 0.75.

More intuitively, the skew-symmetry condition enforces negative pairwise feedback across the community. For example, if species *j* promotes the growth of species *i* (ie. *J*_*ij*_ *>* 0), and species *i* reduces the growth of species *j* (*J*_*ji*_ ≈ −*J*_*ij*_), then a small increase in species *j* will boost species *i*, which feeds back to check the initial growth of species *j*. Similarly, with skew-symmetric off-diagonal elements in the Jacobian, no pair of species simultaneously harms or benefits one another, preventing positive feedbacks that cause the system to diverge even further from the equilibrium when perturbed. **In summary of our fourth main result, specific correlations between the pairwise and higher-order interactions, and within the higher-order interaction network itself, allow stability even when the underlying system cannot be stable with pairwise interactions alone**.

Higher-order interactions that generate skew-symmetric net effects between all pairs of species promote coexistence, but there are also simpler ways for higher-order interactions to have the same effect. One straightforward example is to choose 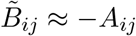 for the inter-specific interactions, meaning that the summed higher-order effects on species *i* involving species *j* are of the same magnitude but opposite sign of the pairwise effect of species *j*. In this way, the net interactions in the Jacobian are all approximately zero (see Fig. G of the Supplementary Information). Indeed, higher-order interactions that effectively turn off the net interactions between species have been found to stabilize diverse coexistence in previous work on microbial interactions [15].

In sum, our results suggest that there are many ways to create stable target abundance equilibria through specific choices of the higher-order interactions. Higher-order interactions that precisely counter the pairwise interactions and those that create skew-symmetric patterns are likely just two examples of a broader class of stabilizing structures. Using a simple stochastic optimization algorithm, we found many other possible stabilizing patterns (see Fig. I of the Supplementary Information), but a large portion of them had structures that were similar to the skew-symmetric patterns we illustrate in Fig. 4.

## Discussion

Reconciling diverse coexistence in nature with our detailed understanding of species interactions has proved to be a remarkably difficult problem, stimulating research in theoretical and empirical ecology for decades [1, 2, 3, 6]. Coexistence does not easily emerge from models with only pairwise interactions, whether these interactions are sampled from probability distributions [6, 23] or measured experimentally [7]. On the other hand, diverse communities in nature are commonplace, from the thousands of coexisting bacterial strains in the soil [33] to the hundreds of tree species in the Amazon rainforest [34, 35]. One possible resolution to the tension between pairwise models and empirical coexistence is that interactions among three or more species are one of the main drivers of diversity in nature. Here, we imposed this scenario on a theoretical model by ensuring that higher-order interactions balanced the pairwise interactions that would otherwise exclude species. We then determined when this procedure permitted robust coexistence and quantified the patterns in the higher-order interactions that succeeded in doing so.

We found that even when higher-order interactions are set exactly to generate a given set of abundances, and thus enable a coexistence equilibrium while the pairwise interactions favor exclusion, the equilibrium need not be stable. Alone, this result challenges efforts to explain species coexistence with higher-order interactions, because even the “right” interactions might only generate unstable equilibria. Moreover, we showed that competitive pairwise systems in which facilitative higher-order interactions enabled a coexistence equilibrium were less likely to be stable than facilitative pairwise systems in which competitive higher-order interactions generated the coexistence equilibrium. Given that many of the natural systems for which coexistence mechanisms remain unresolved are known to be structured by competitive pairwise interactions, this result throws up a further challenge to coexistence via higher-order interactions. In short, the facilitative higher-order interaction solution is an inherently unstable one. We also showed that regardless of the sign of the pairwise or higher-order interaction, the weaker the higher-order interaction strength, the more stable the resulting equilibrium. For similar reasons, setting species target abundances further from where the pairwise interactions take the system, and therefore requiring stronger higher-order interactions, made for less stable equilibria.

Why is it that, even when higher-order interactions are constrained to generate coexistence, they often still fail to do so? The feasibility of the target abundance equilibrium is determined by the sum of the pairwise and higher-order interactions, yet local stability is determined by the pairwise partial derivatives. The constraints on the higher-order interactions ensure that the total effect of the interactions affecting each species is small, but in doing so, they still produce net interactions between each pair of species that eventually destabilize the equilibrium. However, this does not mean that higher-order interactions cannot generate stable coexistence. This outcome simply requires a higher-order interaction network that also makes the net effects of each species on each other either small or have a specific stabilizing structure. In our last set of results, we showed that correlations between the pairwise and higher-order interactions which produced these types of relationships could indeed stabilize diverse coexistence.

This result mirrors previous results for pairwise interactions alone [23], which showed that negative correlations between specific pairwise species interactions promoted stability. In this context, it should not be surprising that higher-order interactions that either entirely negated the pairwise interactions or produced net interactions with strong negative feedback gave rise to stable communities. However, because these higher-order correlation structures are finely tuned to the underlying pairwise interactions, we do not expect them to emerge in every ecological community.

On the other hand, correlations between the pairwise and higher order interactions may be perfectly reasonable in some biological situations or with specific mechanisms of interaction. Previous work has shown that higher-order interactions among the production and counteraction of antibiotics can produce coexistence [15, 19]. Another potential mechanism for stabilizing higher-order interactions is cross-feeding [36, 37, 38], in which microbial species consume preferred nutrients and secrete byproducts that can be utilized by other taxa. Because metabolite production networks have a tree-like structure [39, 40], species that specialize on resources at the root of the tree mostly suffer from competitive interactions, while species that principally grow on byproducts benefit from emergent, multi-species nutrient exchange. As a result, this combination of pairwise resource competition and higher-order nutrient exchange could produce the sorts of stabilizing interaction structures we have identified here.

Our first set of results demonstrated the challenges of maintaining coexistence through randomly sampled higher-order interactions, consistent with recent theory for higher-order interactions in generic ecological models [20]. On the other hand, correlations between the pairwise and higher-order interactions maintained diverse coexistence, recovering the qualitative results of previous models focused on specific biological situations [15, 16]. Our work therefore demonstrates that a coupling between the pairwise and higher-order interactions is a key determinant of community stability, unifying previous results for how higher-order interactions impact coexistence.

There are of course many potential explanations for the mismatch between measured pairwise interactions and observed coexistence that do not involve higher-order interactions [41]. Spatial structure, environmental heterogeneity or poorly constructed pairwise models could all lead to the same phenomenon. In this light, direct empirical measurements of the higher-order interactions may be necessary to clarify whether or not they are responsible for coexistence in a given system [17, 42, 43], and our theory provides specific patterns that can be searched for in these data. More generally, our results suggest that higher-order interactions need to have a precise relationship with highly variable pairwise interactions if they are to stabilize diverse coexistence. These sorts of relationships would essentially never appear by chance, suggesting that specific biological mechanisms are necessary to generate them. An important direction for future work, therefore, is to explore whether or not mechanistic models of species interactions naturally give rise to the pairwise and higher-order interaction structures that stabilize coexistence.

## Methods

### Sampling procedure

Although the mean and between-species variability of the higher-order interactions are set by the pairwise interactions and the target abundances, the variability in the higher-order interactions that affect a given species is unconstrained. Here, we sample the higher-order interaction parameters uniformly from the space of all possible solutions where the higher-order interactions have the required sign. To compute constrained higher-order interactions for a given set of pairwise interactions *A*_*ij*_ and equal target abundances 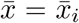, we sample *S* times from a Dirichlet distribution with 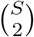 categories and equal concentration parameters. We then multiply each of these 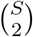 values by the appropriate sum of growth rates and interactions ie. 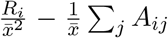. Then, the higher-order interactions modifying species *i*’s growth sum to the correct quantity (ie. 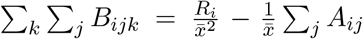). In short, the higher-order interactions are completely determined by the specified pairwise interactions, the target abundances and our subsequent constraint-based sampling procedure. As a result, we can investigate how coexistence in these models is affected by changing only the mean and variance of the pairwise interactions. We analyze alternative sampling methods in the Supplementary Information, where we vary the within-species variability of the higher-order interactions as another free parameter.

When the target abundances are variable between species, this simple sampling scheme no longer works because weighted sums of the higher-order interaction parameters are required. We employ a “Hit and Run” sampling method to generate the required samples [44, 45] (see the Supplementary Information for more detail and the associated code in the GitHub repository at https://github.com/theogibbs/CoexistenceHOIs). To construct matrices that illustrate our skew-symmetric Jacobian in Fig. 4, we sample from a anti-correlated bivariate normal distribution and adjust its row sums to determine the matrix *J* and then compute the appropriate 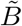 matrix from the formula 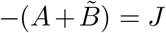. Note that, in this case, we have not actually specified all of the higher-order interactions (instead only specific sums of them) so the higher-order interactions that modify the growth of a given species may have differing signs, unlike in our usual sampling procedure. To solve the dynamics of Equation 1 throughout this work, we used the Livermore Solver for Ordinary Differential Equations (LSODA) from the deSolve v1.25 package [46] in R version 3.6.1.

### Jacobian and stability calculations

The (*i, j*)-th entry of the Jacobian of the model in Equation 1 evaluated at the target abundance equilibrium is

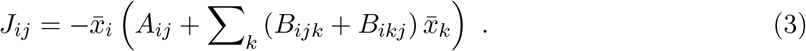

To determine whether or not a target abundance equilibrium is stable, we compute the eigenvalues of this matrix and ask whether or not the the real parts of these eigenvalues are negative. We now appeal to results from ecological random matrix theory [47] to predict when this system becomes unstable. We expect one eigenvalue of this matrix to be determined by the mean interaction strength [47]. Specifically, it will be given by the average row sum of *J* and have a corresponding eigenvector with all equal entries. When the target abundances are the carrying capacities, the average row sum is

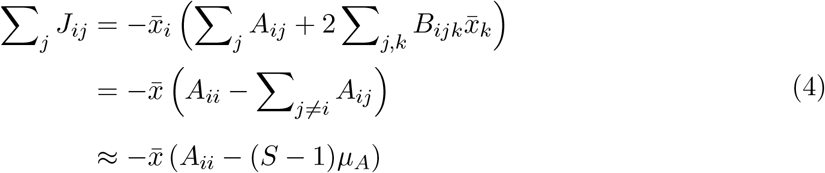

where we have used the constraints (Equation 2) in the second equality. For a system with purely pairwise interactions, the analogous eigenvalue is given by the formula 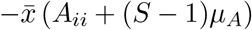. As we describe in the main text (and show numerically in Fig. D of the Supplementary Information), the only difference between these formula is the sign of the mean pairwise interaction strength *μ*_*A*_. Turning to the effect of the variation in interaction strengths on the eigenvalues, we expect the entry-wise variance of the Jacobian to control the radius of a circular distribution of the eigenvalues in the complex plane. This result is called the circular law (see [47, 48, 49] and Fig. A of the Supplementary Information). Let’s take 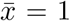, and neglect the correlations between the pairwise interactions and the higher-order interactions, as well as between the higher-order interactions themselves. Then, the variance of a single entry of the Jacobian is approximately

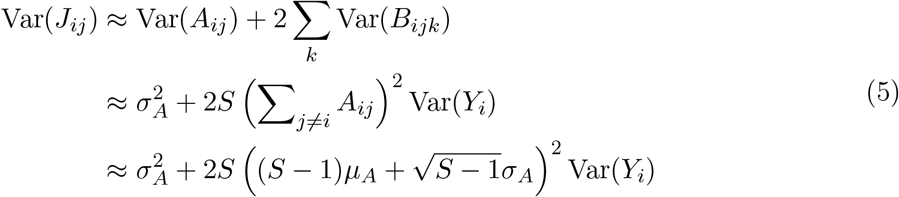

where *Y*_*i*_ is the *i*-th sample from a symmetric Dirichlet distribution with 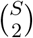 trials and we have used the constraints (Equation 2) in the second line. We have also approximated the average value of the sum of the inter-specific coefficients as 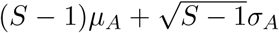. to account for the fluctuations around this value arising from the variable interactions. The Dirichlet distribution has variance 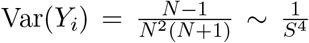 where 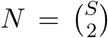. As a result, the entry-wise variance scales like 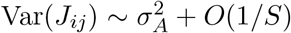 with the number of species *S*, suggesting that the variation in pairwise interactions (or the mean pairwise interaction strength from the previous analysis) controls stability. This is a heuristic analysis and does not comprise a mathematical proof of our results, though it does reproduce the dependence we observe in simulations. In the Supplementary Information, we carry out a similar analysis of the case when there are *S* = 3 species.

## Supplementary Information

We provide additional calculations and simulation results.

## Acknowledgements

We thank members of the Levin and Levine labs for helpful comments and discussion. T.G. was supported by the National Science Foundation Graduate Research Fellowship Program under Grant No. DGE-2039656. Any opinions, findings, and conclusions or recommendations expressed in this material are those of the author(s) and do not necessarily reflect the views of the National Science Foundation. T.G. and J.M.L. acknowledge support from NSF Grant DEB-2022213. S.A.L. acknowledges support from NSF Grant DMS-1951358. K.S.M. acknowledges support from NSERC Discovery 400353.

