## Supplementary Information for "Can higher-order interactions resolve the species coexistence paradox?"

#### Contents


1 Feasibility and stability in ecological models 2

2 Three species calculation 2

3 Example spectra 3

4 Variable growth rates 4

5 Alternative sampling procedures 4

6 Disentangling feasibility and stability 5

7 The effect of target abundances 5

8 Correlated pairwise interactions 6

9 Correlations between pairwise and higher-order interactions 7

10 Bifurcations and initial conditions 7

11 Figures 8

See the GitHub repository at https://github.com/theogibbs/CoexistenceHOIs for all the code to run simulations and generate figures.

### 1 Feasibility and stability in ecological models

For all $S$ species to coexist at equilibrium in Equation 1 of the main text, two conditions have to be met. First, there must be positive abundances ($x_i^\star$) for which each of the $S$ differential equations is zero – a condition known as feasibility [30]. Ecologically, the feasibility condition first ensures that all species are present and then measures whether or not there is any species that could grow to larger or decrease to smaller abundance. Second, the densities must return to this equilibrium after small perturbations, which is known as stability. Feasible abundances are stable if all of the eigenvalues of the Jacobian evaluated at this equilibrium have negative real part. Intuitively, stability measures the system's ability to rebound after small changes in the abundances. If a system is unstable, even tiny changes in species abundances can send the community to an entirely different composition.

### 2 Three species calculation

When there are only three species, there are just three inter-specific higher-order interactions (HOI) and it is straightforward to solve for them using the constraints in Equation 2 of the main text. Additionally, these are the unique higher-order interactions that give rise to the target abundances. The higher-order interactions are

$$B_{ijk} = \frac{1}{\bar{x}_j \bar{x}_k} \left( R_i - \sum_{l=1}^{3} A_{ij} \bar{x}_l \right) = \frac{R_i}{\bar{x}^2} - \frac{1}{\bar{x}} \sum_{l=1}^{3} A_{ij} \quad \text{(S1)}$$

where the second equality is only true if every species has the same target abundance. With these interactions, the Jacobian becomes

$$J_{ij}(\bar{x}) = \begin{cases} -R_i + \bar{x} \sum_{k \neq j} A_{ik}, & i \neq j \\ -\bar{x} A_{ii}, & i = j \end{cases} \quad \text{(S2)}$$

so we can express the stability properties of the constrained higher-order interaction case purely as a function of the pairwise interactions. In fact, we can further simplify this formula by noting $\bar{x} \sum_{k \neq j} A_{ik} = \bar{x}\left(A_{ii} + A_{ik}\right) = R_i + \bar{x} A_{ik}$ where $k \neq i, j$ when the target abundances are the carrying capacities $\bar{x} = R_i/A_{ii}$. Therefore, we find that the Jacobian is

$$J_{ij}(\bar{x} = R_i/A_{ii}) = \begin{cases} \bar{x}A_{ik} \text{ where } k \neq i, j, & i \neq j \\ -\bar{x}A_{ii}, & i = j \end{cases} \tag{S3}$$

It is instructive to compare this formula with the Jacobian of the purely pairwise model with the same equilibrium abundances $\bar{x}$, which is given by $J_{ij} = -\bar{x}A_{ij}$. There are two differences. First, each off-diagonal entry of the HOI Jacobian (Equation S3), does not have a negative sign. Second, these off-diagonal entries are permuted with respect to the pairwise Jacobian, so that the inter-specific interactions within each row have been swapped.

Basic results from ecological random matrix theory [47] yield predictions for how the statistics of the interactions affect stability. Though this theory is only formally justified in diverse communities (the $S \to \infty$ limit), we find that it also describes the behavior of less diverse communities over many replicates. The mean interaction strength determines one eigenvalue of each Jacobian, whose corresponding eigenvector has approximately the same value across the entries [47]. As a result, this eigenvalue will be approximately given by the average row sum of the Jacobian. In the pairwise model, this average row sum is $-\bar{x}((S-1)\mu_A + \mu_D)$ where $\mu_D$ is the average self-regulation coefficient ($A_{ii}$) and $\mu_A$ is the average off-diagonal pairwise interaction coefficient. On the other hand, the row sum eigenvalue of the constrained higher-order Jacobian is given by $\bar{x}\left((S-1)\mu_A - \mu_D\right)$ because of the difference in sign of the off-diagonal elements. These formula explain the results observed in Fig. 2 of the main text, where competitive pairwise interactions destabilized the equilibrium obtained via constrained higher-order interactions more readily than mutualistic pairwise interactions. They are also consistent with our calculation for a diverse community in the Methods. Because neither of the modifications (the sign or the swapping) affect the overall variability of the Jacobian, Equation S3 also shows that variation in the pairwise interactions produces the same quantitative stability properties for models with and without constrained higher-order interactions. For the parameterizations we have considered, the swapped entries in Equation S3 do not affect stability, but this can occur when there are additional correlations between the pairwise interaction coefficients (see the section titled Correlated pairwise interactions).

### 3 Example spectra

The spectra of the Jacobian evaluated at the target abundance equilibrium depend on both the variation in the interactions and the mean interaction strength (Fig. A). To understand how these statistics affect the eigenvalue distribution, we plot spectra from the constrained hgiher-order case and the case with only pairwise interactions where feasibility is ensured through growth rates (see the Disentangling feasibility and stability section for more details). For both of these cases, one eigenvalue is determined by the mean interaction strength, but its sign has differing dependencies on the mean pairwise interaction strength (Fig. A and [47]). Specifically, competitive pairwise interactions ($\mu_A > 0$) produce a negative real eigenvalue for the feasible pairwise case, but a positive real eigenvalue for the constrained higher-order case (Fig. A), and vice versa for facilitative pairwise interactions. These eigenvalues produce the stability dependence we observed in Fig. 2 of the main text. Similarly, when the pairwise interactions are highly variable, this variability controls the probability of stability by setting the radius of the circular distribution of eigenvalues (Fig. A and [47]). Interestingly, when the variability in pairwise interactions is smaller and the mean interactions strength is non-zero, the variability in higher-order interactions controls the radius of the eigenvalue distribution (Fig. A). In the Disentangling feasibility and stability section, we investigate exactly how stability depends on these different statistics more completely for both of these cases.

#### 4 Variable growth rates

In the main text, we set each growth rate to the same value $R_i = 1$, but these can instead vary between the species. Intuitively, we expect this variation to further decrease the probability of stability because it makes the constrained higher-order interactions more variable as well through the constraint equations:

$$\sum_k \sum_j B_{ijk}\bar{x}_j\bar{x}_k = R_i - \sum_j A_{ij}\bar{x}_j \ . \tag{S4}$$

Simulations show that the probability of stability decreases as a function of the variability in growth rates when they are sampled from normal distributions with increasing variance (Fig. B). In fact, variable growth rates can eventually produce instability even when there are no inter-specific pairwise interactions imply because the constrained higher-order interactions must become large and variable to produce feasibility.

#### 5 Alternative sampling procedures

Throughout our results in the main text, we assumed that the constrained higher-order interactions affecting a given species have the sign required to produce the desired target abundance. We then sample uniformly from the space of all possible solutions to this problem. If we allow each higher-order interaction to have arbitrary sign, however, then the space of possible solutions is no longer a set with finite volume, so the constraint problem is vastly under-determined. Therefore, we can introduce a new free parameter – the variability in the constrained higher-order interactions ($\sigma_B$) – which will also affect stability (Fig. C). Specifically, we can add additional noise (with variance $\sigma_B^2$)

to the constrained higher-order interactions we found using our usual algorithm while ensuring the constraint equations remain true. Increasing the variation in the constrained higher-order interactions decreases the probability of stability (Fig. C), as we would expect. The sampling procedure we employ in the main text is more likely to be stable than constrained higher-order interactions with additional noise, but the variation in the constrained higher-order interactions from our usual sampling method still affects the spectra (Fig. A), so it is not obvious exactly how these constrained higher-order interactions affect stability overall.

#### 6 Disentangling feasibility and stability

In the standard Lotka-Volterra model (ie. Equation 1 of the main text without higher-order terms), we can choose the growth rates to precisely balance the pairwise interactions ($R_i = \bar{x} \sum_j A_{ij}$), implying that the target abundances are feasible [23]. We then construct an analogous higher-order interaction model with the same pairwise interactions but where feasibility is instead guaranteed through the constraints on the higher-order interactions. As we described in the main text, these two models display identical stability properties (Fig. D). In the first set of simulations, we set the mean interaction strength to zero and specify a small amount of variability in the interactions. We then progressively increase the number of species in the system to observe when it becomes unstable. A classic result due to May [6] and others [23] shows that larger communities with pairwise interactions are less likely to be stable. Here, we observe quantitatively the same behavior for constrained higher-order interactions (Fig D). Results for non-zero mean interaction strengths are analogous, except for the reversed dependence on the mean interaction strength we described in our first result (Fig D). In short, constrained higher-order interactions are not generating an additional stability benefit beyond guaranteeing feasibility, as predicted by our characterization of the eigenvalues in the Methods section of the main text.

#### 7 The effect of target abundances

In the main text, we showed that stability had an interesting dependence on the value of the target abundances, where some target abundances were more stable than others. In fact, there appears to be a specific target abundance value for each set of interactions that maximize stability (Fig. EA). Conversely, the statistics of the pairwise interactions also change which value of the target abundances is most stable (ie. the minima of the curves in Fig. EA). More competitive pairwise interactions produce smaller stable target abundances, and vice versa (the different lines in Fig. EB). Interestingly, larger variation in the pairwise interactions produces larger stable target abundances for any average strength of the pairwise interactions (ie. the curves in Fig. EB all increase). Overall, however, increased variation in the pairwise interactions always eventually leads to instability (Fig. EC), even when the target abundances are tuned to the underlying pairwise interactions. In other words, even though some target abundances were more stabilizing than others, no choice of target abundances could stabilize a community with sufficiently variable pairwise interactions.

To sample from the space of constrained higher-order interactions with variable abundances, we used the R package hitandrun [44, 45]. When the abundances are all equal, the higher-order interactions are sampled uniformly from a large dimensional simplex (ie. from a Dirichlet distribution), but variable abundances introduce a bias in this sampling space (see Fig. L for a low dimensional example). To compute the most stable abundances in Fig. E, we used the R function optimize from the package stats, while ensuring the optimization did not hit the boundary conditions on the search.

#### 8 Correlated pairwise interactions

In the Three Species Calculation section, we compared the Jacobia of the feasible pairwise and constrained higher-order models in the case when there were only three species. Besides a sign difference, the Jacobia had permuted dependence on the pairwise interactions, suggesting that structure in the pairwise interactions that affected stability in the feasible pairwise case would have less of an effect on the higher-order case. Simulations of more diverse communities show that this is indeed the case (Fig. F). Specifically, we impose a correlation between off-diagonal entries of the pairwise interactions (ie. $A_{ij}$ and $A_{ji}$). In the feasible pairwise case, positive correlations decrease the probability of stability while negative correlations increase it [23]. In fact, a perfect negative correlation guarantees stability (see [23] and Fig. F). Interestingly, the same qualitative dependence holds for constrained higher-order interactions, with more negative pairwise correlations increasing the probability of stability and vice versa. The correspondence is not quantitatively the same, however, and perfect negative correlations that assured stability in the feasible pairwise case no longer do so for constrained higher-order interactions. This is because the permuted Jacobian of the constrained higher-order case does not have the same finely tuned structure as a system controlled by feasible pairwise interactions alone.

#### 9 Correlations between pairwise and higher-order interactions

In the main text, we noted that tight coupling between the pairwise interactions $A_{ij}$ and all the higher-order interactions that modify this pairwise interaction $\tilde{B}_{ij}$ can essentially guarantee stability by producing nearly diagonal Jacobia. Fig. G displays one such example of this phenomena, where this relationship between pairwise and higher-order interctions (ie. $A_{ij} \approx -\tilde{B}_{ij}$) produces a Jacobian with eigenvalues clustered around the self-regulation parameters: $-A_{ii} = -1$.

To discover other possible stabilizing higher-order interaction structures, we ran a simple stochastic optimization algorithm on the higher-order interactions for a given system with pairwise interactions. At each step of the optimization, we randomly permuted the higher-order interactions that affect each species (therefore maintaining equality in the constraint equations) and also added mean zero random noise to all of the parameters. When the real part of the leading eigenvalue (the eigenvalue with largest real part) decreased on account of these modifications, we accepted these new higher-order interactions. Running this algorithm over many steps usually produced spectra of the target abundance equilibrium with non-zero imaginary parts and real parts concentrated at the self-regulation parameters (Fig. H). Interestingly, this is exactly the form of the spectra that we expect from imposing skew-symmetry on the Jacobian itself. Although the resulting Jacobia are certainly not precisely skew-symmetric, they are considerably more negatively correlated across the diagonal than the inital Jacobia from randomly sampled constrained higher-order interactions (Fig. I).

#### 10 Bifurcations and initial conditions

Throughout this work, we have determined when the target abundance equilibrium is stable, but we have not investigated the dynamics of Equation 1 from the main text when this equilibrium is unstable. Simulations of these unstable systems reveal a wide range of possible behaviors, including species exclusion, limit cycles, complex fluctuations and diverging abundances. Interestingly, there are new feasible and stable equilibria in this regime – a situation that can never arise in a Lotka-Volterra model with only pairwise interactions. As the mean pairwise interaction strength becomes more competitive, the target abundance equilibrium eventually becomes unstable, at which point the dynamics are actually attracted to a second stable equilibrium (Fig. JA). In fact, the second equilibrium becomes unstable at the same value of the mean interaction strength as the pairwise model with guaranteed feasibility, significantly broadening the range of mean pairwise interaction strengths in which constrained higher-order interactions produce coexistence (Fig. JB). Importantly, the equilibria in these models are not necessarily globally stable, and the size of the basin of attraction can vary with the interaction statistics themselves (Fig. K). A more complete investigation of all the equilibria of models with structured higher-order interactions, as well as their basins of attraction, is an important direction for future work.

#### 11 Figures

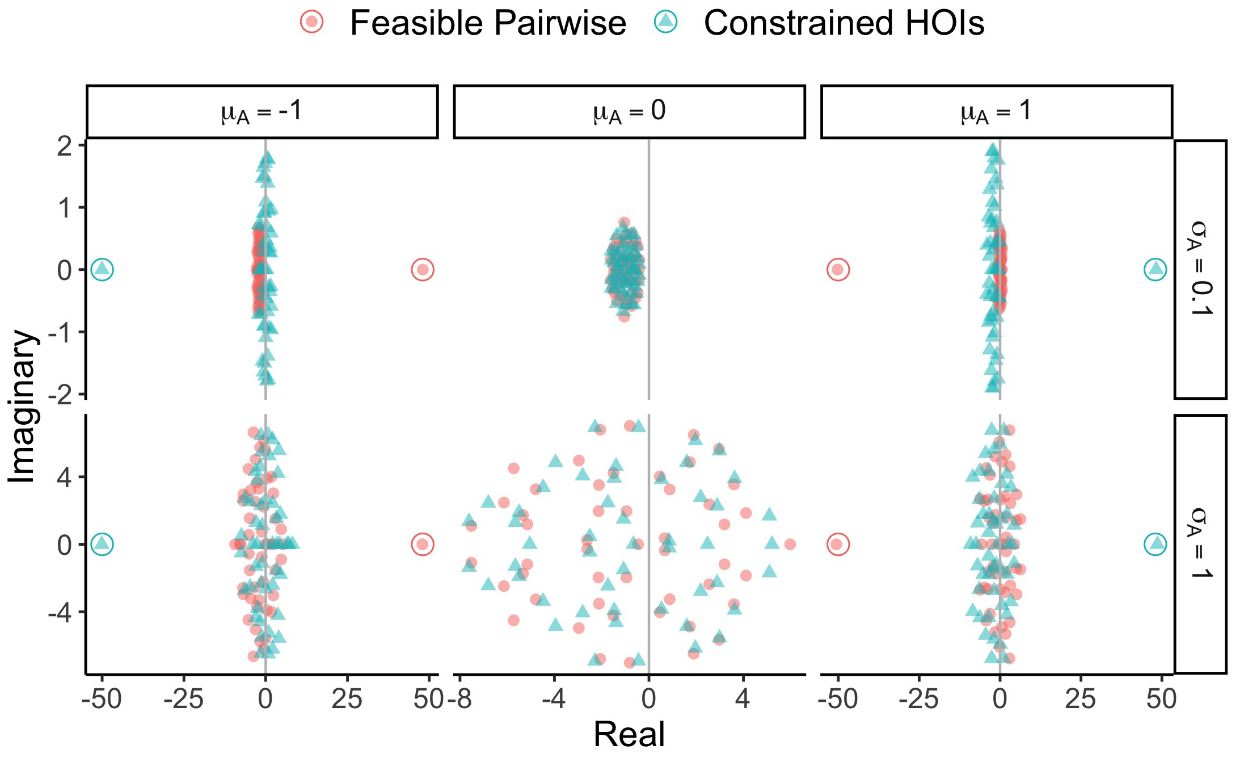


Figure A: The distribution of eigenvalues in the complex plane for Jacobia from the feasible pairwise or constrained higher-order models (colors and shapes) evaluated at the target abundance equilibrium. Different columns designate different mean interaction strengths while different rows show different variations in pairwise interactions. There are $S = 50$ species. Gray vertical lines show the imaginary axis and colored circles display our predictions for the one eigenvalue determined by the mean interactions strength.

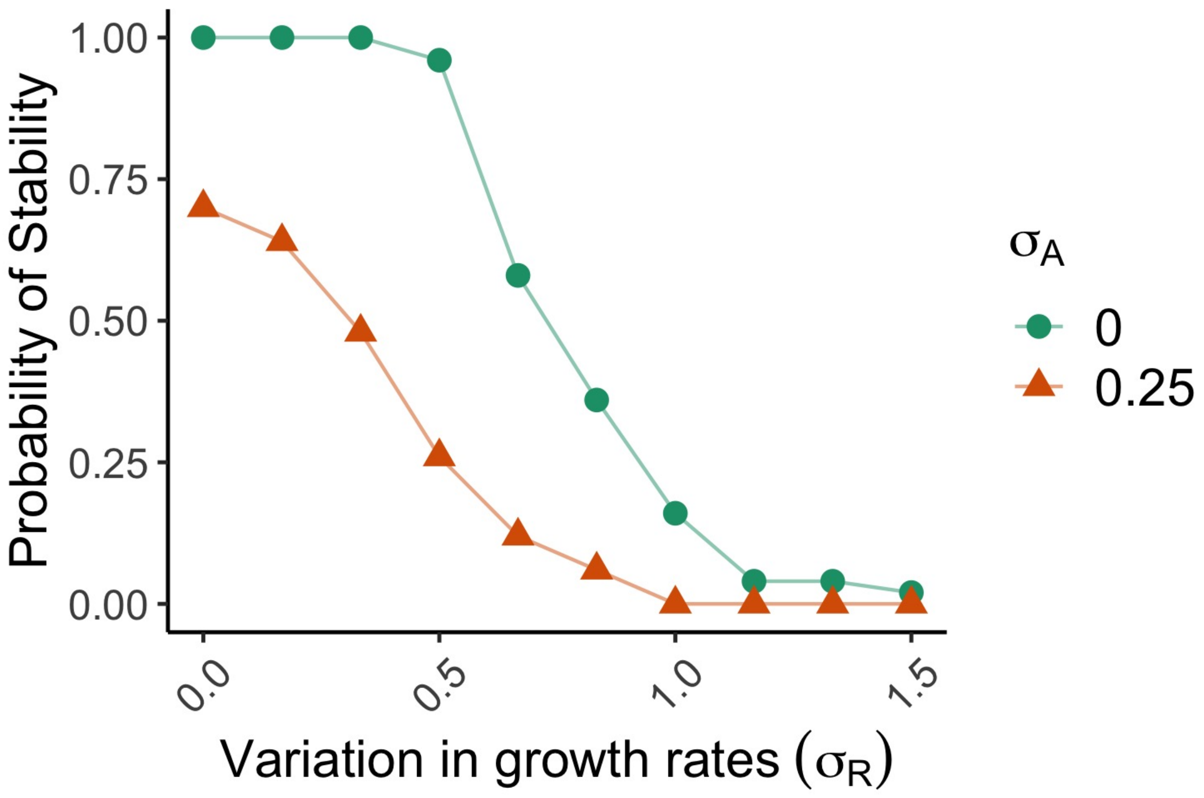


Figure B: The probability of stability as a function of the variation in growth rates (the standard deviation $\sigma_R$ of a normal distribution with mean $1$). The different colors and shapes designate different variations in pairwise interactions. The probabilities are computed over $50$ replicates, there are $S = 20$ species and $\mu_A = 0$.

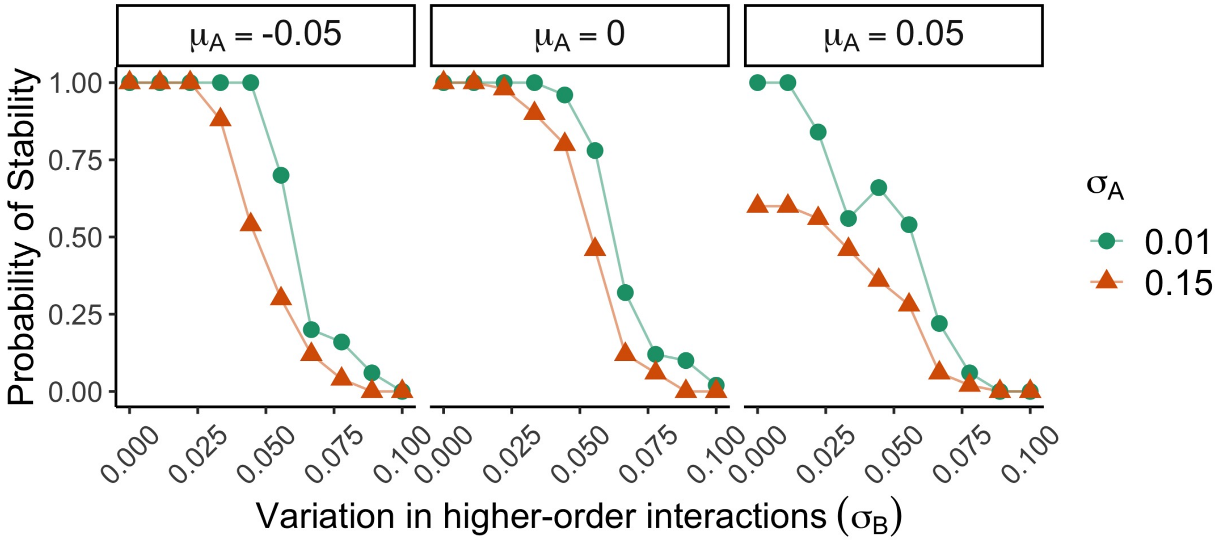


Figure C: The probability of stability as a function of the variation in constrained higher-order interactions (the standard deviation $\sigma_B$ of a normal distribution with mean precisely zero added to higher-order interactions that satisfy the constraints). Columns designate different values of the mean interaction strength while colors and shapes designate different variations in pairwise interactions. Probabilities are computed over $50$ replicates and there are $S = 20$ species.

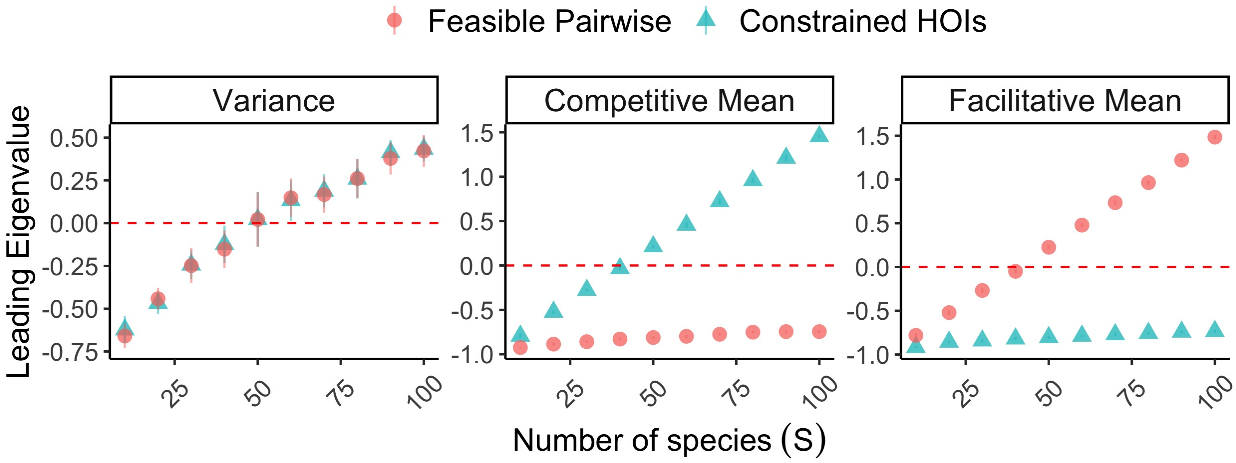


Figure D: (Variance Panel) We plot the real part of the leading eigenvalue (the eigenvalue with largest real part) as a function of the number of species. Shapes show the average and error bars show $\pm 1$ standard deviation over $10$ replicates. The red dashed line denotes the stability threshold at zero. For each point, the mean pairwise interaction strength is $\mu_A = 0$ and the variation in pairwise interactions is $\sigma_A = 0.15$. (Competitive Mean Panel) The same as in the Variance panel but with $\sigma_A = 0.025$ and $\mu_A = 0.025$. (Facilitative Mean Panel) The same as in the Variance panel but with $\sigma_A = 0.025$ and $\mu_A = -0.025$.

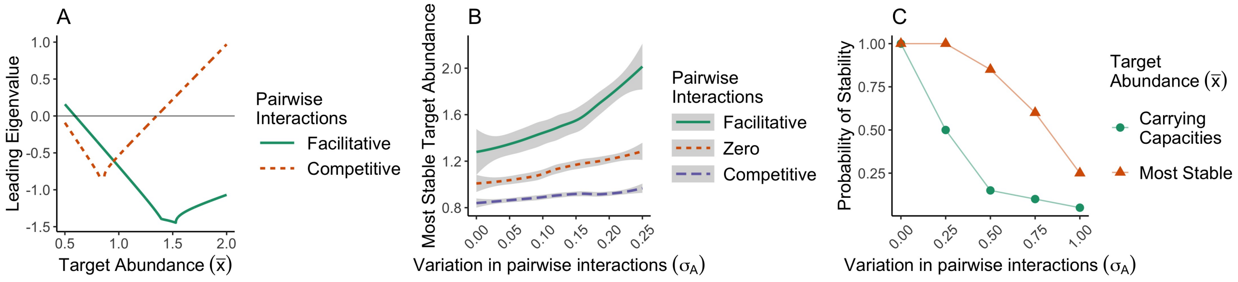


Figure E: (A) The real part of the leading eigenvalue (the eigenvalue with largest real part) as a function of the common target abundance ($\bar{x}$). The different colors and linetypes designate differing values of the mean pairwise interaction strength – facilitative denotes $\mu_A = -0.25$ and competitive denotes $\mu_A = 0.25$. Each of the different curves uses the same random draw of interactions, explaining their similar geometries. There are $S = 3$ species and the variation in pairwise interactions is $\sigma_A = 0.25$. (B) The average (lines) and standard error (shaded areas) of the most stable target abundance value (the minima in Fig. 3B of the main text) as a function of the variation in pairwise interactions. The colors and linetypes designate different values of the mean pairwise interaction strength – facilitative denotes $\mu_A = -0.1$, zero denotes $\mu_A = 0$ and competitive denotes $\mu_A = 0.1$. There are $S = 5$ species and the averages are computed over $5$ replicates of $50$ different variations in pairwise interaction strengths. (C) The probability that the target abundance equilibrium is stable as a function of the variation in the pairwise interactions. The colors and shapes denote whether or not the target abundances are set to the carrying capacities of the species or the most stable target abundance. There are $S = 5$ species and $\mu_A = 0$.

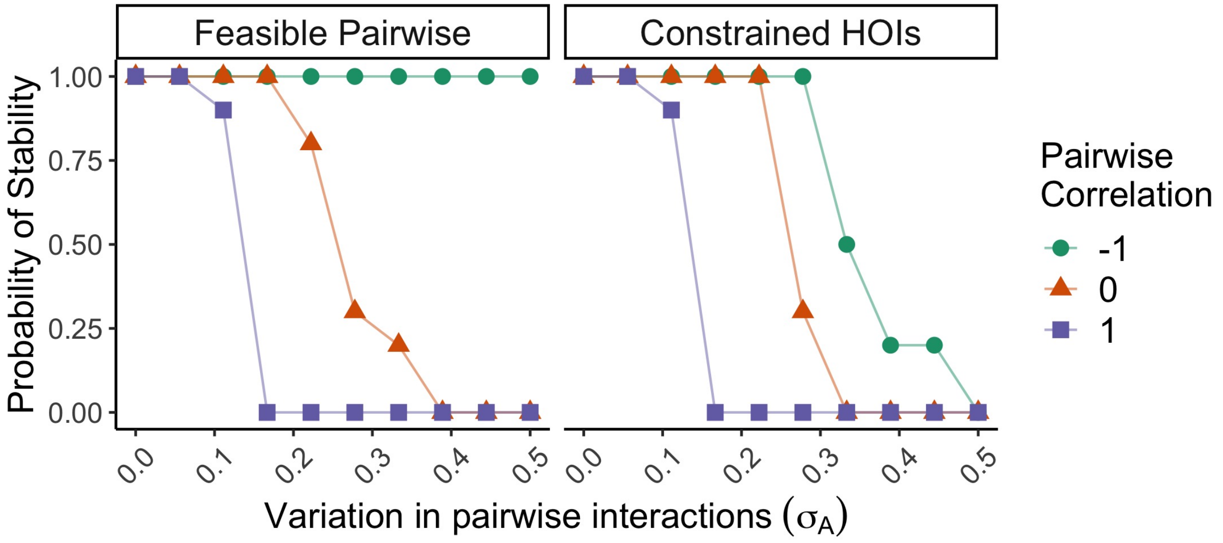


Figure F: The probability of stability as a function of the variation in pairwise interactions for the feasible pairwise and constrained higher-order models (columns) for different values of the correlation between off-diagonal pairwise interactions (colors and shapes). Probabilities are computed over $10$ replicates and there are $S = 20$ species.

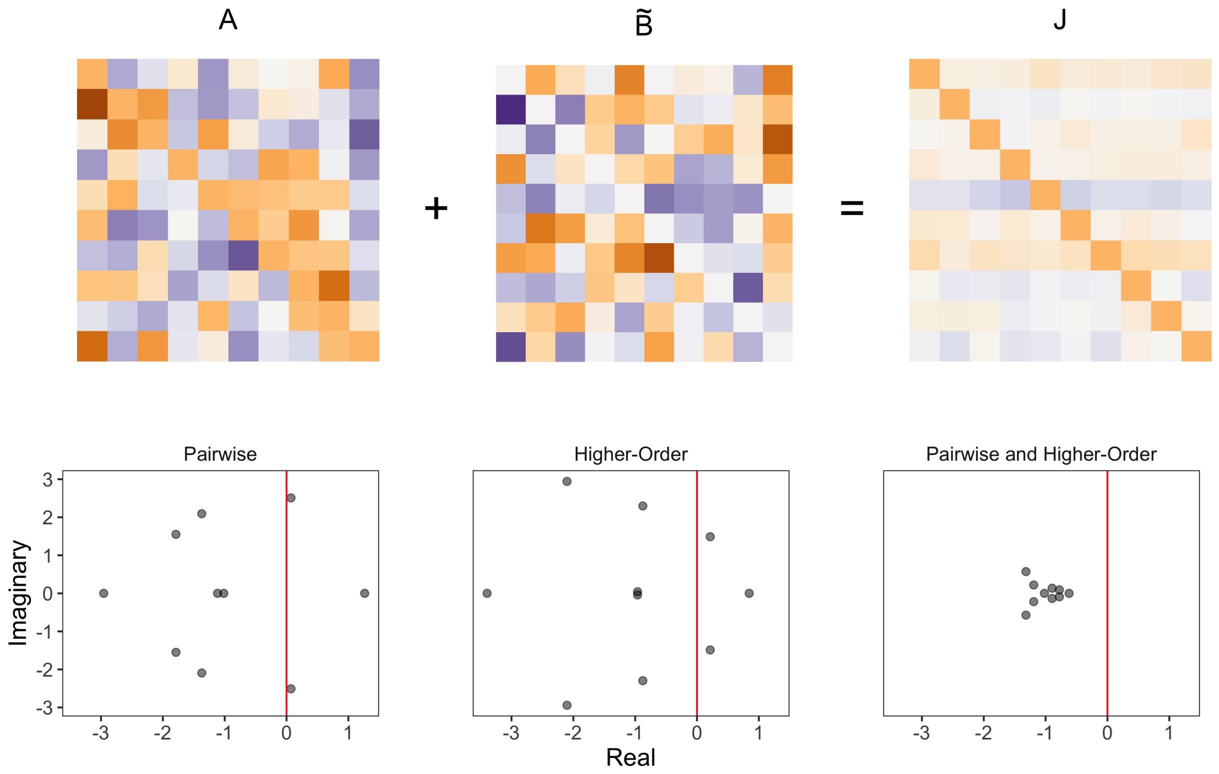


Figure G: The top row shows the pairwise interaction matrix $A$, the matrix collecting higher-order interactions $\tilde{B}$ and the Jacobian which is a sum of the two matrices. Darker orange colors indicate more negative values, while darker purple colors indicate more positive values. The bottom row shows the spectra of the system with only pairwise interactions, only higher-order interactions and both types of interactions. There are $S = 10$ species, $\mu_A = 0$ and $\sigma_A = 1$. See https://github.com/theogibbs/CoexistenceHOIs for exact details on how the $\tilde{B}$ matrix is generated.

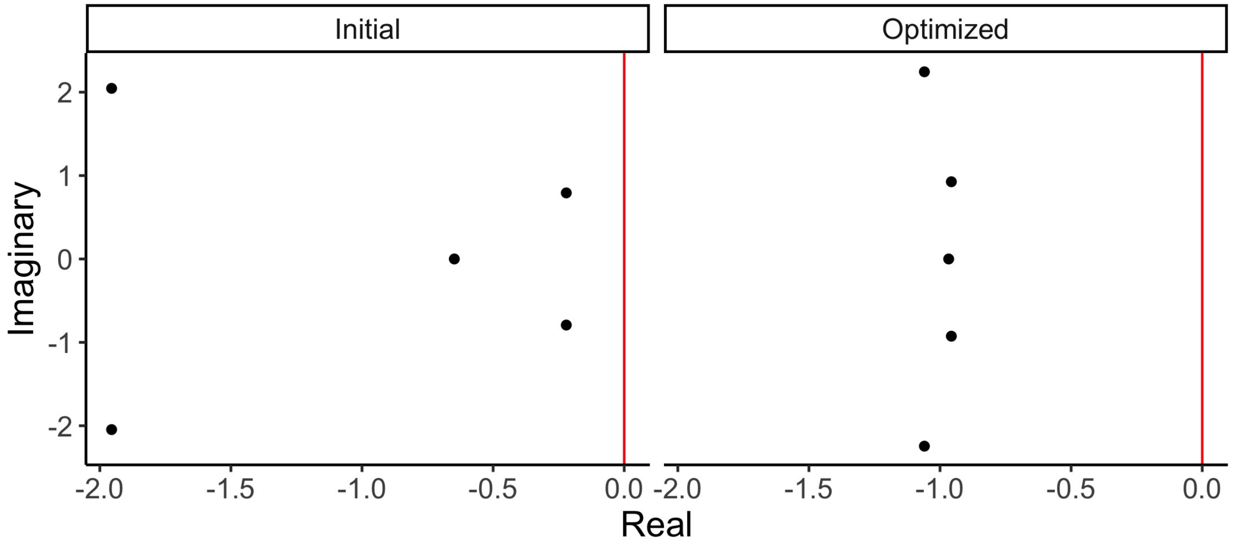


Figure H: The spectra of the Jacobian before the optimization on the higher-order interactions (labeled Initial) and afterwards (labeled Optimized). There are $S = 5$ species, $\mu_A = 0$ and $\sigma_A = 0.5$.

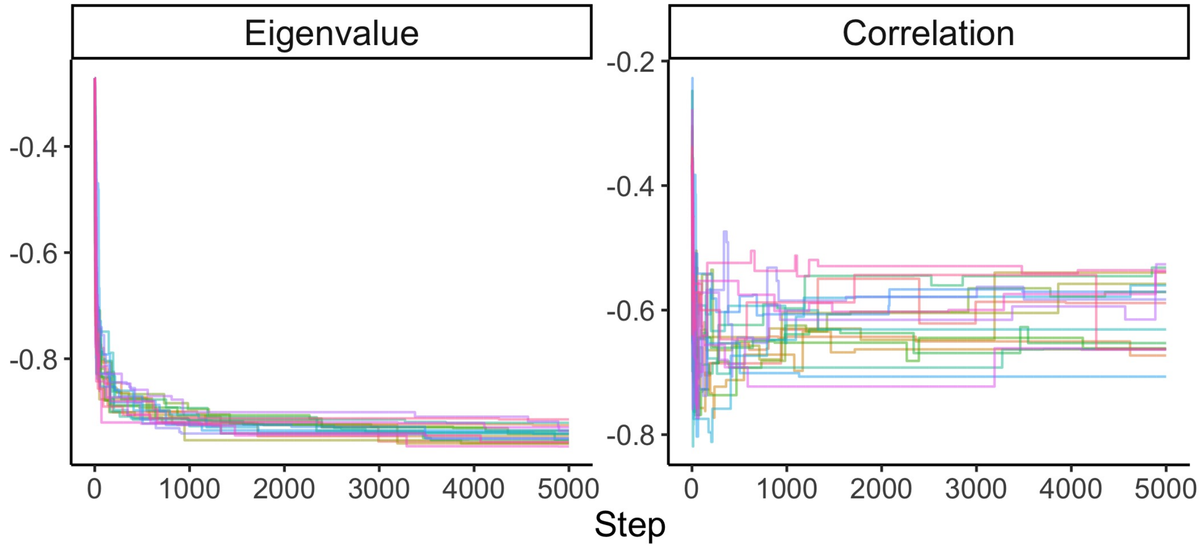


Figure I: The real part of the leading eigenvalue (labeled Eigenvalue) and the correlation between off-diagonal elements of the Jacobian (labeled Correlation) as a function of the current step in the optimiziation. The different colors designate different replicates of $20$ total. All cases use the same underlying pairwise interactions with $S = 5$ species, $\mu_A = 0$ and $\sigma_A = 0.5$.

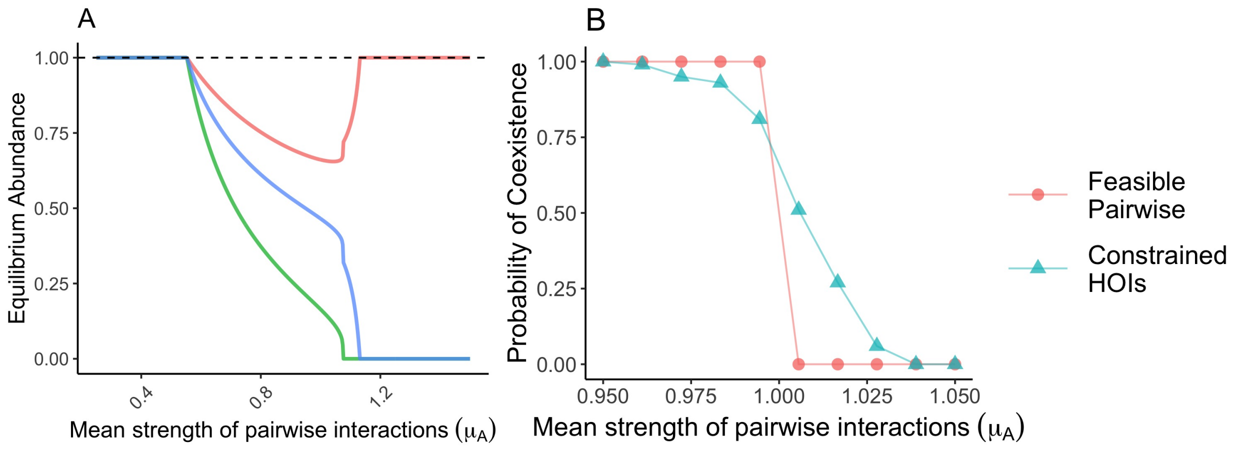


Figure J: (A) We plot the equilibrium abundances at the end of a simulation of the dynamics as a function of the mean strength of the pairwise interactions. Different colors designate different species. The dashed vertical line shows the value of the target abundances. Initially, the species stably coexist at these target abundances but this equilibrium becomes unstable and they coexist at different abundances until two species are eventually excluded. (B) The probability that all species have positive abundance at the end of a simulation of the dynamics as a function of the mean strength of the pairwise interactions. Different shapes and colors denote whether or not the model has feasible pairwise interactions or constrained higher-order interactions. There are $S = 20$ species and no variation in the pairwise interactions $\sigma_A = 0$.

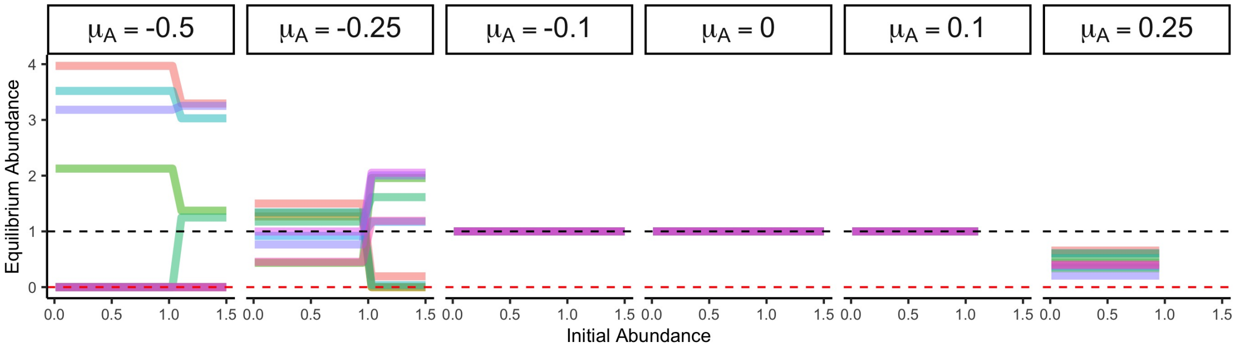


Figure K: The equilibrium abundances at the end of a simulation of the dynamics as a function of the initial starting abundance common to all species. Colors designate different species and columns show different mean interaction strengths. We set the variance in interaction strengths to $\mu_A/\sqrt{3}$ to match the statistics of a uniform distribution. The black dashed line shows the target abundance equilibrium, while the red dashed line shows zero abundance. There are $S = 10$ species. When there is no line to show a solution, the abundances quickly diverge so our numerical procedure does not produce a solution. For weak interactions that are not variable, the abundances converge to the target abundance equilibrium regardless of the initial conditions. When the interactions are stronger, other equilibria can be reached, including one where all species coexist (as in Fig. J). Similarly, the dynamics may blow up and not reach an equilibrium.

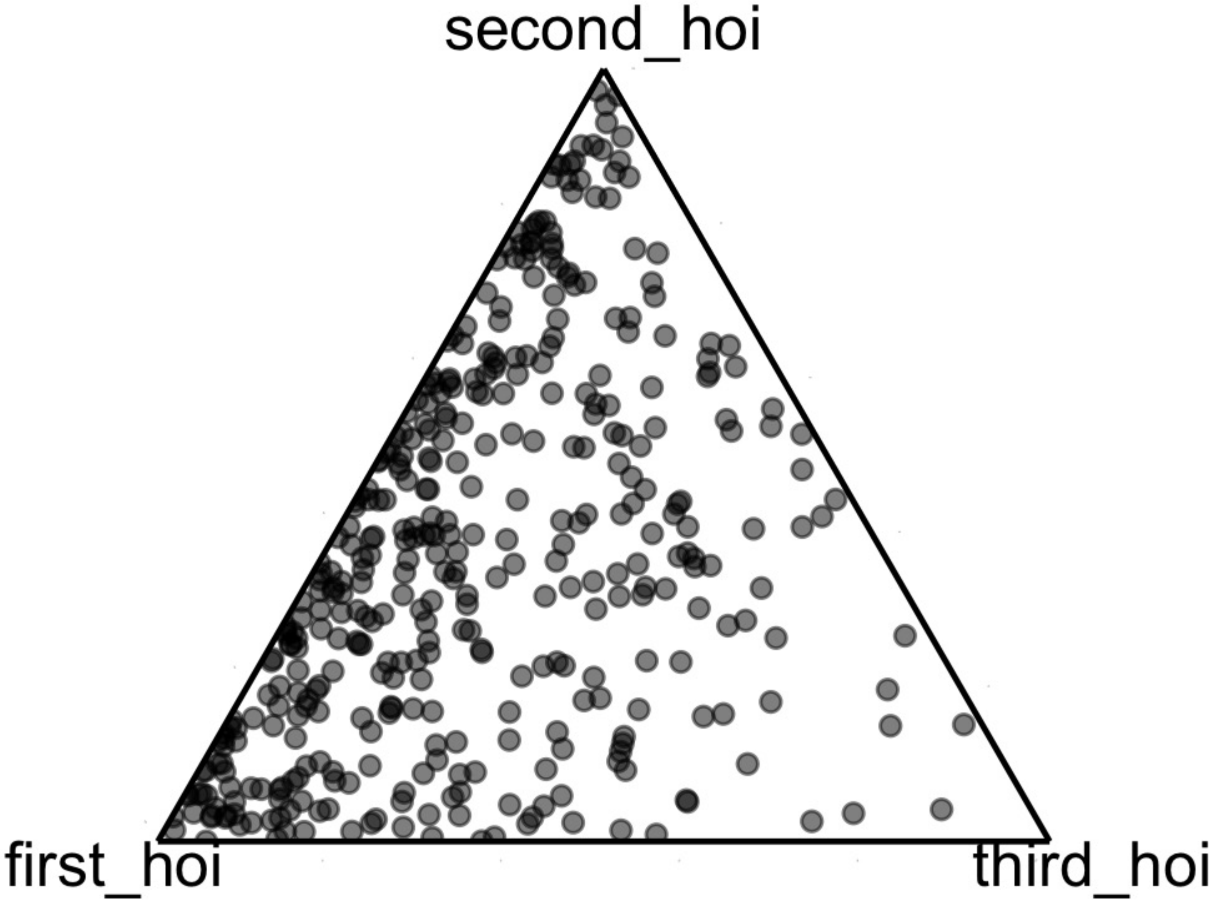


Figure L: A simplex (or ternary) plot showing the bias in higher-order interactions produced by variable abundances over $100$ replicates for each of the species. There are $S = 4$ species so each species experiences $3$ higher-order interactions, meaning that they can be plotted easily. The variation in pairwise interactions is $\sigma_A = 0.5$ and the target abundances are $x_1 = 0.1$, $x_2 = 0.2$, $x_3 = 0.9$ and $x_4 = 0.8$. Higher-order interactions involving these larger abundances must be smaller to maintain equality in the constraints, producing the visible leftward bias. In the case of equal abundances, the higher-order interactions would be uniformly distributed on this simplex.
